# The impact of ER^UPR^ on mitochondrial integrity mediated by PDK4

**DOI:** 10.1101/2024.07.12.603217

**Authors:** Priyanka Mallick, Sebabrata Maity, Rupsha Mondal, Trina Roy, Oishee Chakrabarti, Saikat Chakrabarti

## Abstract

ER and mitochondrial stress are often interconnected and considered as major contributors to aging as well as neurodegeneration. Co-ordinated induction of ER^UPR^ and mito^UPR^ has been observed in diabetes and pulmonary disorders. However, in the context of aging and neurodegeneration, regulation of this intra-organellar crosstalk has remained relatively elusive. Here, we demonstrate that pyruvate dehydrogenase kinase 4 (PDK4), a mitochondrial protein accumulates at the ER-mitochondrial contact sites (MAMs) during ER stress. Classically, PDK4 is known to phosphorylate PDHA1 (pyruvate dehydrogenase E1 subunit alpha 1), and plays a significant role in regulating the oxidative phosphorylation driven ATP production. In this study, we propose a non-canonical function of PDK4; we show that it acts as a connecting link between ER^UPR^ and mito^UPR^ with significance in aging and neurodegeneration. Transcriptomics analyses show increased PDK4 levels upon drug induced ER stress. We detect elevated PDK4 levels in brain lysates of Alzheimer’s disease (AD) patients, as well as in *in vivo* and *ex vivo* AD models. Additionally, exogenous expression of PDK4 was found to refine ER-mitochondrial communication, significantly altering mitochondrial morphology and function. Further, we also observe defective autophagic clearance of mitochondria under such conditions. It is prudent to suggest that elevated PDK4 levels could be one of the key factors connecting ER^UPR^ with mito^UPR^, a phenotypic contributor in aging and at least some neurodegenerative diseases.

## Introduction

Endoplasmic reticulum (ER) plays a central role in the folding process of membrane and secretory proteins; proteins that fail to attain their proper conformation due to misfolding, unfolding or erroneous modifications get cleared off from the cell. However, under certain physiological stresses, like nutrient deprivation, import-export or storage imbalance of Ca^2+^, aging or pathological insults, like bacterial or viral infections, neurodegeneration, to name a few, the folding capacity at the ER gets compromised. This leads to accumulation of misfolded proteins within the ER lumen, termed as ER stress, and elicits a cellular response known as the unfolded protein response (UPR) ^1^. Recent studies suggest that ER stress in various cellular contexts can alter mitochondrial morphology and bioenergetics through transcription dependent and independent pathways ^2–5^.

Mitochondria are indispensable cellular organelles that regulate ATP generation, Ca^2+^ signaling ^6,7^, apoptosis ^6,8^, reactive oxygen species (ROS) production and neutralization ^9^, neurotransmitter metabolism ^10^ etc. Thus, maintaining mitochondrial homeostasis is extremely crucial for cellular homeostasis and viability. In neurons, spatio-temporal regulation of the mitochondrial functions is attained with mitochondrial localization ^11, 12^ and biogenesis ^13^, processes that are largely dependent on the fission-fusion competence of these organelles and their transport efficiency. Hence, perturbations in mitochondrial dynamics, bioenergetics and altered activity of their fission-fusion machineries are observed in most neurodegenerative diseases and neuropathies ^5, 14–16^. However, the contribution of altered mitochondrial dynamics in disease etiology is still far from being fully understood. While significant efforts have been put in studying the contributions of the canonical ER^UPR^ pathways (ATF6, IRE1α and PERK) in the context of ER stress mediated neuronal disease progression, identification and functional characterization of components from the other organelles, especially the mitochondria, remains to be studied.

ER forms membrane contact sites (MCS) with different organelles and plasma membrane. Among such contact sites, ER-mitochondria tethering points or MAM (mitochondria-associated endoplasmic reticulum membrane) junctions are vital in regulating Ca^2+^, metabolites, and lipid transfer between the two organelles ^17^. Thereby, structural integrity and lipid composition of mitochondrial membrane are largely dependent upon imported lipids or precursor molecules from ER ^18^. Further, previous studies have shown that autophagosomes could originate at these ER-mitochondria contact sites under starvation ^19^. These junctions are also suggested to be critical in regulating reticulo-mito-phagy ^20^. Atypical lipid trafficking between ER and mitochondria, as well as defects in autophagy are implicated in ageing and neurodegenerative diseases, supporting the hypothesis that ER^UPR^ can directly affect mito^UPR^, a stress response pathway that provides mitochondrial quality control. Increasing evidences show that the number, duration and dynamics of molecular composition of MAM junctions are vital to mitigate cellular stress and neurodegenerative disease pathology ^21–23^. However, detailed understanding of the molecular players in this is warranted.

The pyruvate dehydrogenase complex (PDHc) provides the link between glycolysis and the TCA cycle by catalyzing the end product of glycolysis, pyruvate into acetyl-CoA ^24^. The activity of the PDHc is negatively regulated by its reaction products acetyl-CoA and NADH, and pyruvate dehydrogenase kinase (PDK) dependent serine phosphorylation on the α-subunit (E1) ^24–26^. Pyruvate dehydrogenase phosphatase (PDP) can dephosphorylate and re-activate PDHc ^27^. PDK family of proteins comprise 4 members, PDK1 through 4; all of these have tissue specific expression and activity ^28^. PDKs play significant roles in maintaining the metabolic status of cells. Apart from their function in cellular metabolism, one of the PDKs, pyruvate dehydrogenase kinase 4 (PDK4) has been reported to localize at the ER-mito contact sites of muscle cells, regulating the insulin signaling during obesity ^29^, suggesting an unexplored role for this mitochondrial protein in cellular physiology. Here, we show that eliciting ER^UPR^ can affect PDK4 levels, an enrichment very significantly detected at the ER-mitochondria contact sites. Such an alteration in PDK4 protein is physiologically relevant as its increased levels are detected in serum starved cells. Hippocampi of 1 year old rat brains as well as Alzheimer’s disease (AD) patient derived whole brain lysates corroborate these results. While ER stress elicits higher PDK4 levels in cellular systems, the reciprocal is also true – increasing PDK4 protein levels, upregulates well-established ER stress markers, along with those of mito^UPR^. Elevated PDK4 levels at MAM junctions eventuate to mitochondrial fragmentation and compromised metabolism, enhanced commencement of autophagy but impaired clearance of autophagosomes. Our results unravel non-canonical activity of PDK4 in maintaining mitochondrial structure, function, and quality control; all factors in turn responsible for overall cellular homeostasis and dysregulation of which are often linked with aging and neurodegeneration.

## Results

### Effect of ER stress on PDK4

To induce ER stress, human neuroblastoma SH-SY5Y cells were treated for 3 hrs with 0.5µM thapsigargin (TG), a known chemical inhibitor of sarco endoplasmic reticulum Ca^2+^ ATPase (SERCA) pumps. mRNA transcriptomes of DMSO (control) and TG (drug) treated cells were compared to identify the deregulated genes (log_2_FC ± 2.0 and p-value 0.01) affected by ER stress (Figure 1A). These genes were further mapped onto the MitoCarta database version 2.0 ^30^ to identify mitochondrial genes that were altered during ER stress, using the human specific dataset. Our analyses showed significant upregulation in the mRNA level of the mitochondrial Pyruvate dehydrogenase kinase 4 (PDK4) under ER stress induced by TG treatment (Figure 1B). This upregulation of PDK4 was further confirmed by RT-PCR and Western blot analyses (Figures 1C-1E). mRNA and protein levels of the canonical ER stress markers were elevated under similar conditions (Figure 1F, Supplementary Figures 1A, 1B). Increased protein levels of PDK4 were also detected upon treatment with 5μg/ml tunicamycin (Tun) for 3 hrs, an antibiotic that induces ER stress by abrogating N-linked glycosylation of proteins (Supplementary Figures 1C, 1D) – suggesting a correlation between chemically induced ER stress and PDK4. Together, these results indicate that ER stress can significantly alter the expression of an important mitochondrial kinase, PDK4.

**Figure 1:**
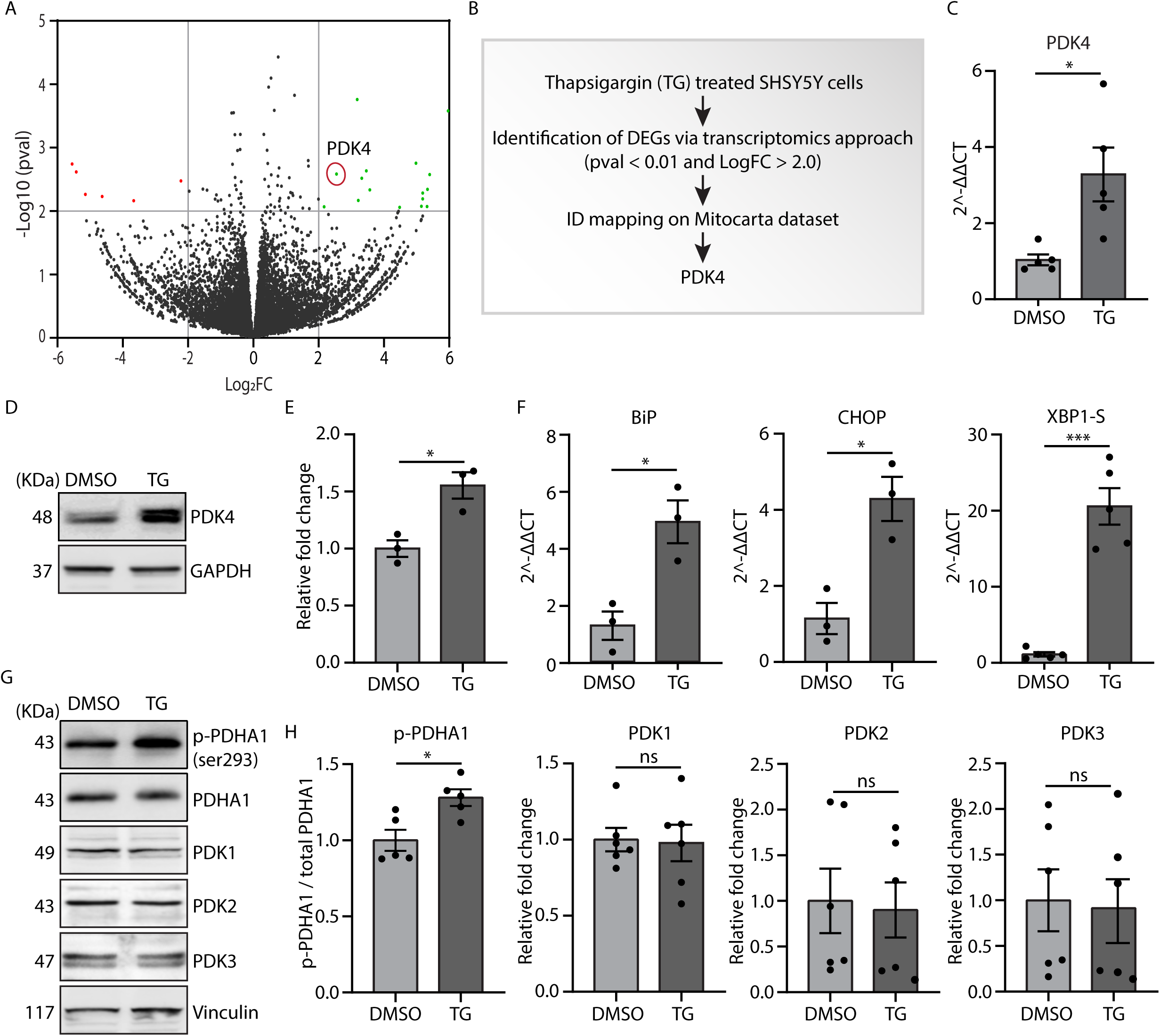
ER stress induces PDK4 expression. (A) SH-SY5Y cells were treated with TG (Thapsigargin, 0.5µM for 3 hrs) or DMSO, and were subjected to RNA transcriptomics analysis. Green and red data points on the volcano plot represent the up and down regulated genes (LogFC ± 2.0 and p-value < 0.01 cut offs marked in grey lines), obtained from three biological replicates of DMSO and TG treated samples, respectively. (B) Flowchart for identification of PDK4 from the deregulated genes. (C) qRT-PCR analysis shows enhanced PDK4 mRNA level in TG treated condition, as compared to the control. Data represent the mean ± SEM of 5 independent experiments. (D) Immunoblotting of cell lysates was carried out to check PDK4 protein level in TG induced ER stress. (E) Graph represents the altered PDK4 protein level and represented as the mean ± SEM of 3 independent experiments. (F) qRT-PCR analyses show increased mRNA levels of known ER stress markers (BiP, CHOP, and XBP1-S) in TG induced ER stress, compared to the control. Data represent the mean ± SEM of 3-5 independent experiments. (G) Cell lysates from DMSO and TG treated samples were immunoblotted to detect expression and phosphorylation status of indicated proteins. (H) Graphs represent changes in expression of different proteins, as mentioned in panel G. p-PDHA1 was normalized to total PDHA1 level. Data represent the mean ± SEM of 5-6 independent experiments. *\*p*≤0.05; *\*\*\*p*≤0.001; ns: not significant (estimated via unpaired two-tailed Student’s t-test).

Additionally, a significant increase in phosphorylation of PDK4 substrate PDHA1, at Ser293 was detected in TG treated cells; however, the total PDHA1 level remains marginally less (Figures 1G, 1H). Since, increased phosphorylation of PDHA1 is linked to the deactivation of PDHc; this observation raises the possibility that under ER stress, PDK4 mediated regulatory role might be exclusive of its usual enzymatic function, reported in TCA cycle. To verify if other isoforms of PDK were similarly affected during ER stress, levels of PDK1, PDK2 and PDK3 were compared during ER stress (Figures 1G, 1H). We could not detect any significant change in the protein levels of any of the isoforms, other than PDK4, though all of them shared extensive sequence similarity (Supplementary Figure 1E).

### Altered PDK4 in physiological models of ER stress

Extensive literature suggests a close link between ER stress and a plethora of neuro-pathophysiological conditions ^31–34^. Hence, we hypothesized a correlation between PDK4 and various patho-physiological stresses. To establish this, cells were subjected to serum withdrawal and analysed for the protein levels of PDK4. Our results showed significant upregulation in PDK4 and BiP upon serum starvation (Figures 2A, 2B).

**Figure 2:**
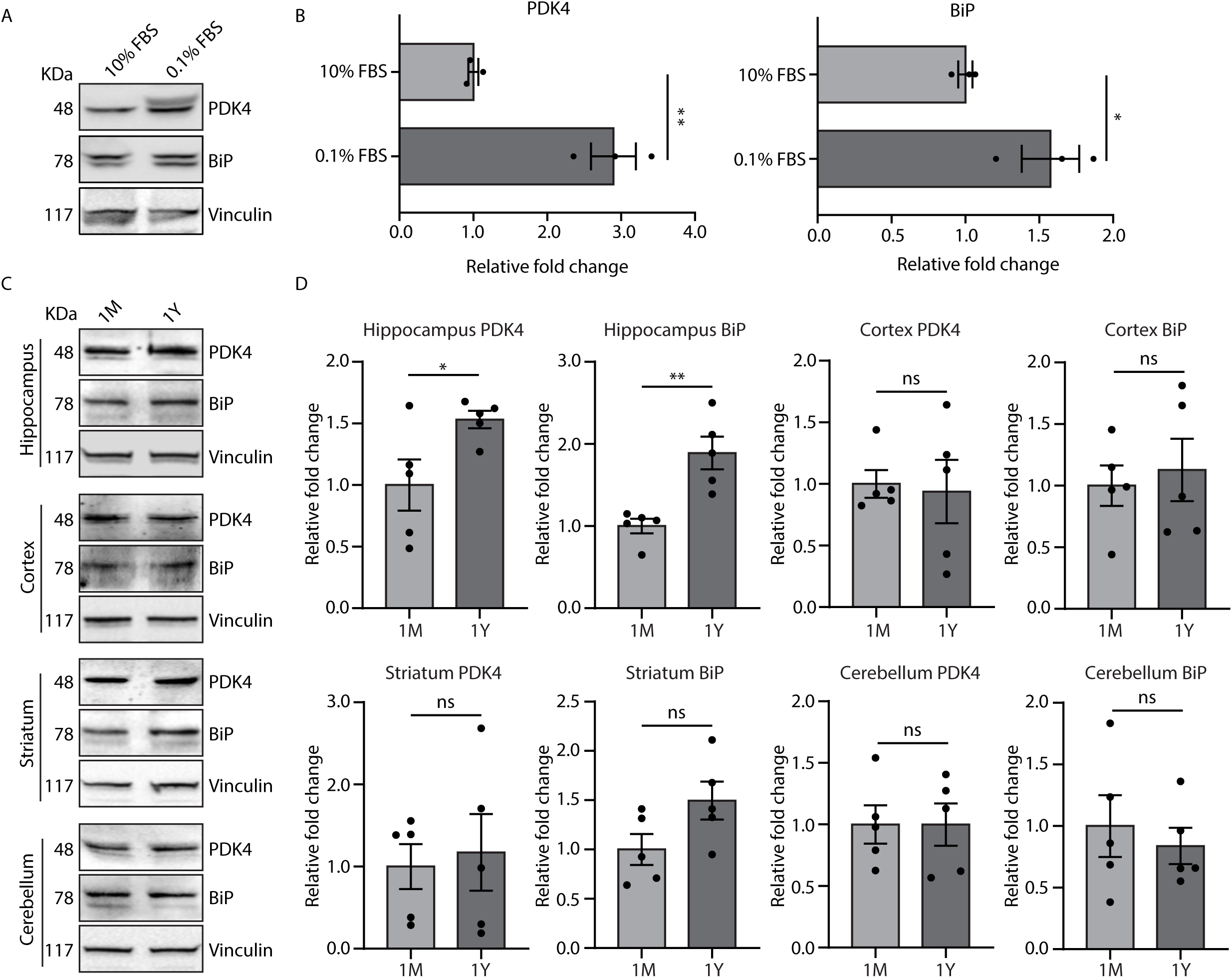
Serum starvation and aging mediated physiological stress elevates PDK4 level. (A) SH-SY5Y cells were treated with highest (10%) and lowest (0.1%) levels of FBS concentrations and immunoblotting of cell lysates were performed to detect PDK4 and BiP protein levels in serum starved condition. (B) Graphs show the alteration in different protein expressions. Data represent the mean ± SEM of 3 independent experiments. (C) Immunoblotting of brain region specific lysates (hippocampus, cortex, striatum, and cerebellum) of 1 month and 1 year old rats were performed, to check the expression of indicated proteins. (D) Graphs represent the altered protein level and represented as the mean ± SEM of 5 animals/group. *\*p*≤0.05; *\*\*p*≤0.01; ns: not significant (estimated via unpaired two-tailed Student’s t-test).

It is well established that cellular proteostasis gets progressively compromised during the process of aging, which is also a known contributing factor in age-related pathologies, like neurodegeneration. Aging as well as neurodegenerative diseases are further associated with ER stress ^35–37^. To explore if PDK4 was affected during aging, its protein levels were verified in different brain regions of young (1 month) and aged (1 year) rats (Figures 2C, 2D). We detected significantly elevated PDK4 levels in aged hippocampus, though not in other brain regions (cortex, striatum and cerebellum). This elevation is correlated with the BiP levels in the various brain regions. Altered neurogenesis of hippocampus is prominent in several disease scenarios including Alzheimer’s disease (AD) ^38^. Additionally, impaired UPR of this region is suggested in aged rats ^39^. Hence, region-specific elevation of PDK4 levels in aged rat brain is indicative of its probable involvement in ER stress mediated hippocampal neurodegeneration.

### Increase in PDK4 levels in pathological models of ER stress

We next aimed to investigate the levels of PDK4 in pathologically relevant models. For this, whole brain lysates of AD patient and control individual were compared; PDK4 protein level was seen to be significantly upregulated in AD brain lysate (Figures 3A, 3B). Due to unavailability of brain region specific AD patient samples, we validated region-specific alterations of PDK4 levels in the well-established animal model of Apolipoprotein E (ApoE) KO mice. Here again, we detected significantly elevated PDK4 protein levels in the hippocampal region of ApoE KO mice (Figures 3C, 3D). The other brain regions did not reflect a similar change in PDK4 levels, recapitulating our observations in aging rat samples (Figures 2C, 2D). Secondly, a significant increase in PDK4 levels was observed in cell-based model of AD when compared with the controls (Figures 3E, 3F). For this, SH-SY5Y cells were transiently transfected with AICD (amyloid precursor protein intracellular domain) followed by treatment with amyloid-β 42 (Aβ 42) fragment ^40^.

**Figure 3:**
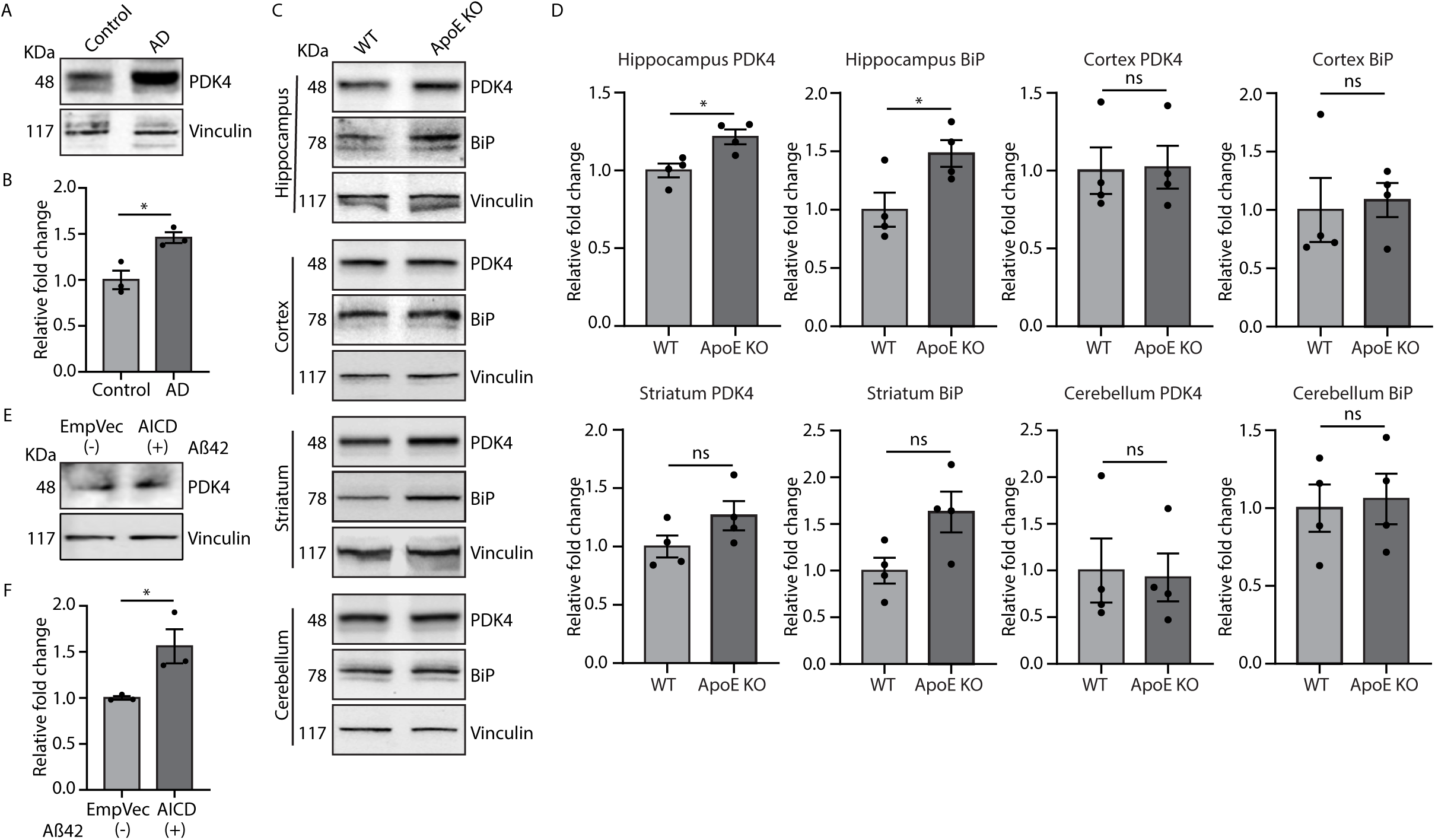
Increased PDK4 expression in Alzheimer’s disease (AD) (A) Whole brain protein lysates of control and AD patients were immunoblotted to check expression of PDK4. (B) Graphs represent the altered PDK4 protein levels and represented as the mean ± SEM of 3 technical replicates. (C) Immunoblotting of brain region specific lysates (hippocampus, cortex, striatum, and cerebellum) of wild type (WT) and ApoE knock out (KO) mice were performed, to check the expression of indicated proteins. (D) Graphs represent the altered protein level and represented as the mean ± SEM of 4 animals/group. (E) SH-SY5Y cells transfected with indicated constructs were treated with DMSO and 0.5CμMCAβ for 48Ch. Cell lysates were immunoblotted to check PDK4 protein level. (F) Graph shows data as the mean ± SEM of 3 independent experiments. *\*p*≤0.05; *\*\*p*≤0.01; ns: not significant (estimated via unpaired two-tailed Student’s t-test).

Thirdly, the status of PDK4 level were analysed in an already characterized prion disease model of neurodegeneration ^41,42^. It has been reported that point-mutations like A117V and KHII (K110I, H111I) can enrich the population of a particular transmembrane form of prion protein (^Ctm^PrP) in cells, a form of PrP associated with sporadic prion diseases. Our results showed that PDK4 was upregulated in presence of the natural Gerstmann-Sträussler-Scheinker syndrome (GSS) variant of PrP (A117VPrP) that is known to generate ^Ctm^PrP in abundance and ER stress (Supplemental Figure 2A, 2B). This suggests a plausible correlation of the presence of higher amounts of PDK4 in multiple neurodegenerative diseases.

Together, these results suggest a plausible role for this mitochondrial protein in the context of ER stress; one of the probable roles could be mediated at the ER-mitochondria junctions.

### PDK4 localizes at MAM junctions during ER stress

An increase in ER-mitochondria junction (or membrane associated mitochondria, MAM) has been previously reported in AD ^43^. In our study, altered presence of PDK4 at the MAM junctions was validated in multiple ways. First, live cell imaging indicated an increase in localization of PDK4 at ER-mitochondria contact sites in cells under ER stress when compared with the control (Figures 4A, 4B); a simultaneous rise in the MAM junctions was also detected (Figure 4C). Secondly, sub-cellular fractionation of cell lysates (Supplementary Figure 3A) showed significantly higher levels of PDK4 protein in MAM junctions in TG treated samples compared to the controls (Figure 4D, 4E). Along with PDK4, other mitochondrial proteins known to be enriched in MAM fractions (MFN2 and VDAC1) were detected at higher levels in TG treated cells than the controls (Figure 4E). Hence, both the imaging and fractionation-based protein estimation results show an increase in MAM junctions and accompanying increase in PDK4 at these contact sites in cells subjected to ER stress. Presence of higher numbers of ER-mitochondria contacts are known to suggest increased mitochondrial fission ^44^ as is seen in cells treated with TG (Supplementary Figures 3B, 3C).

**Figure 4:**
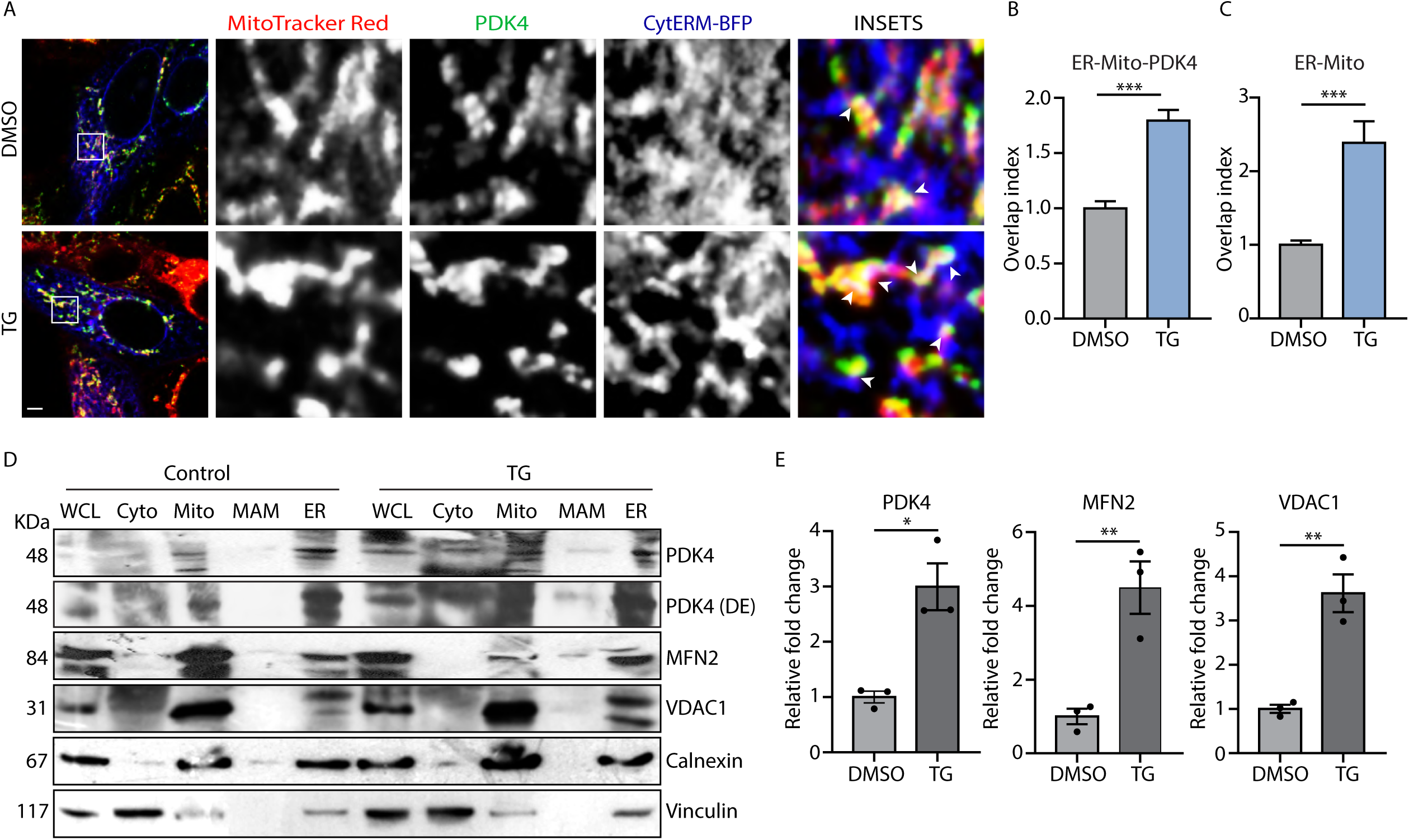
PDK4 localizes at MAM junction under ER stress. (A) SH-SY5Y cells transfected with CytERM–BFP, treated with DMSO or TG (0.5µM for 3 hrs), followed by staining with MitoTracker Red FM. Cells were fixed and immune-stained to detect PDK4 (green). Enlarged views of the areas within the white boxes are shown (insets). Arrowheads mark co-localization of CytERM–BFP (ER), MitoTracker Red (mitochondria) and PDK4 signals (white). Scale bar: 5 µm. (B) Graph shows quantification of CytERM– BFP, MitoTracker Red and PDK4 (as described in A) overlapping; co-localization was represented as mean fold change (overlap index) ± SEM. Data was calculated from ∼60 cells from 3 independent experiments. (C) Graph shows quantification of CytERM–BFP and MitoTracker Red (as described in A) overlapping; co-localization was represented as relative fold change (overlap index). Data was calculated from ∼60 cells from 3 independent experiments. (D) Different cellular fractions obtained from DMSO and TG (0.5µM for 3 hrs) treated SH-SY5Y cells (as described in methods) were subjected to immunoblotting to check the expression of indicated proteins. Both MFN2, VDAC1 were used as Mitochondria and MAM markers; whereas calnexin and vinculin were used as ER and cytosolic markers, respectively. WCL: Whole cell lysate, Cyto: Cytosol, Mito: Mitochondria; DE: Dark exposure. (E) Graphs represent the indicated protein levels and represented as the mean ± SEM of 3 independent experiments. *\*p*≤0.05; *\*\*p*≤0.01; ns: not significant (estimated via unpaired two-tailed Student’s t-test).

### Elevated PDK4 level elicits ER stress-like response

Next, we wanted to verify if increased PDK4 levels could evoke an ER stress-like response and also affect mitochondria. To address this, lysates from SH-SY5Y cells transiently transfected with PDK4 were analysed for known ER^UPR^ marker proteins (Figures 5A, 5B). We observed significantly elevated levels of BiP and ATF5 upon exogenous expression of PDK4; ATF5 is also known to play an important role in mito^UPR^ ^45^. Phosphorylated PDHA1 was enhanced in the PDK4 transfected cells as compared to the control, further confirming intact PDK4 canonical activity. Live cell imaging of PDK4 over-expressed cells showed presence of more MAM junctions in them, when compared with the control (Figures 5C, 5D), though the increase in the number of such contact sites was not as robust as with TG treatment. We also detected an increase in mitochondrial fragmentation in cells with exogenous PDK4 (Figures 5E-5G), confirming changes in mitochondrial morphology upon PDK4 over-expression ^46^. These observations suggest a direct correlation between PDK4, altered mitochondrial morphology and ER stress.

**Figure 5:**
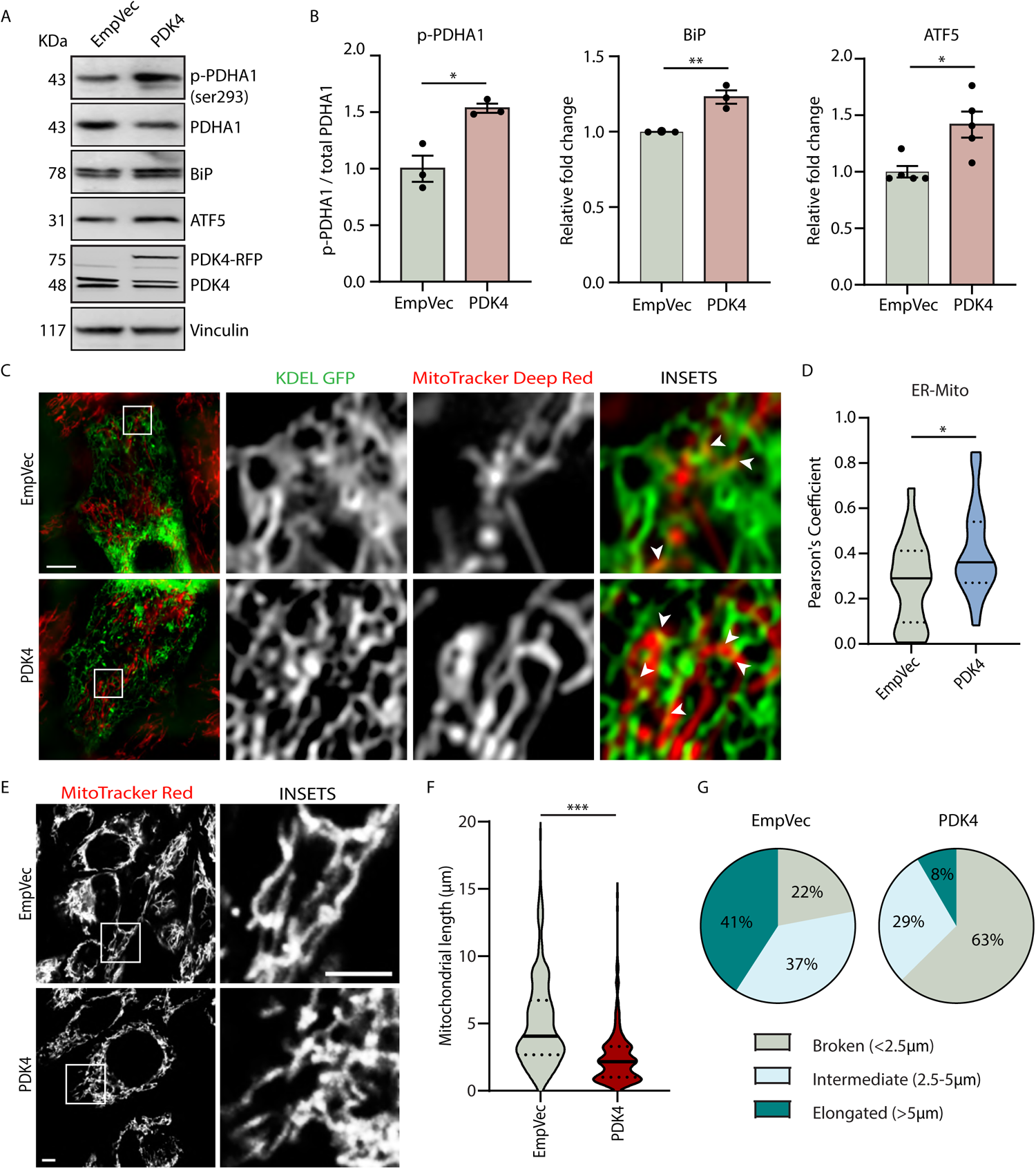
PDK4 over-expression affects mitochondrial morphology and homeostasis. (A) SH-SY5Y cell lysates from empty vector (EmpVec) and PDK4 transfected samples were immunoblotted to detect expression and phosphorylation status of indicated proteins. (B) Graphs represent change in expression of different proteins, as mentioned in A. p-PDHA1 was normalized to total PDHA1 level. Data represent the mean ± SEM of 3-5 independent experiments. (C) Cells transfected with KDEL-GFP, PDK4 and control (EmpVec), imaged under live-cell conditions using MitoTracker Deep Red FM. Enlarged views of the areas within the white boxes are shown (insets). Scale bar: 10 µm. (D) Violin plot shows Pearson’s correlation coefficient for co-localization of KDEL-GFP (ER) and MitoTracker Deep Red (mitochondria) in indicated samples (as described in C). Data shown are of ∼60 cells from 3 independent experiments. Solid and dashed black lines mark median and quartiles respectively. (E) Cells transfected with empty vector (EmpVec) and PDK4, imaged under live-cell conditions using MitoTracker Red FM. Enlarged views of the areas within the white boxes are shown (insets). Scale bar: 5 µm. (F) Violin plot of the mitochondrial length (μm) in images as described in E. Solid and dashed black lines mark median and quartiles respectively. (G) Pie-charts show categorical representation of total mitochondrial pool depending on their lengths (as indicated) in images shown in panel E. *\*p*≤0.05; *\*\*p*≤0.01; ns: not significant (estimated via unpaired two-tailed Student’s t-test).

### Altered PDK4 levels affect mitochondrial function

Not just morphology, increased PDK4 levels also altered mitochondrial function (Figures 6). We observed that exogenous expression of PDK4 led to reduction in total ATP production (Figure 6A). This was most importantly associated with a significant decrease in mitochondrial dependency and pronounced increase in glucose dependency and glycolytic capacity, when compared to the control (Figure 6B). ATP generated from amino acid and fatty acid metabolism was reduced upon PDK4 over-expression. Our results fit well with previous observation that PDK4 mediated phosphorylation and inhibition of PDC complex can negatively affect mitochondria-mediated ATP generation ^47–49^. Next, Western blot analyses of the five different OXPHOS complex subunit proteins (NDUFB8, SDHB, UQCRC2, COX-II, ATP5A for complex I-V respectively) showed significantly low levels of Complex-I protein (NDUFB8) in PDK4 over-expressed condition; proteins of the other complexes remained unaltered across samples (Figures 6C, 6D). However, we could not detect any significant alteration in expression of the representative subunits of OXPHOS complexes (ND2, SDHA, CYTB, COX-II, ATP8 for complex I-V respectively) at the mRNA levels by qRT-PCR (Supplementary Figure 4). Interestingly, flow cytometric analyses with MitoTracker Green and TMRM co-staining suggested an increase in mitochondrial mass in presence of exogenous PDK4 (Figure 6E, 6F). A significantly concomitant decrease in mitochondrial potential was also observed; revealing that the mitochondrial potential to mass ratio was severely compromised in PDK4 over-expressed cells (Figure 6G-6I). These clearly suggest an abundance of dysfunctional (more mass but depolarized) mitochondria. The presence of higher amounts of reactive oxygen species (ROS) was detected using flow cytometric analyses with DCFDA staining, further supported the hypothesis that elevated PDK4 levels promote mitochondrial stress (Figure 6J, 6K).

**Figure 6:**
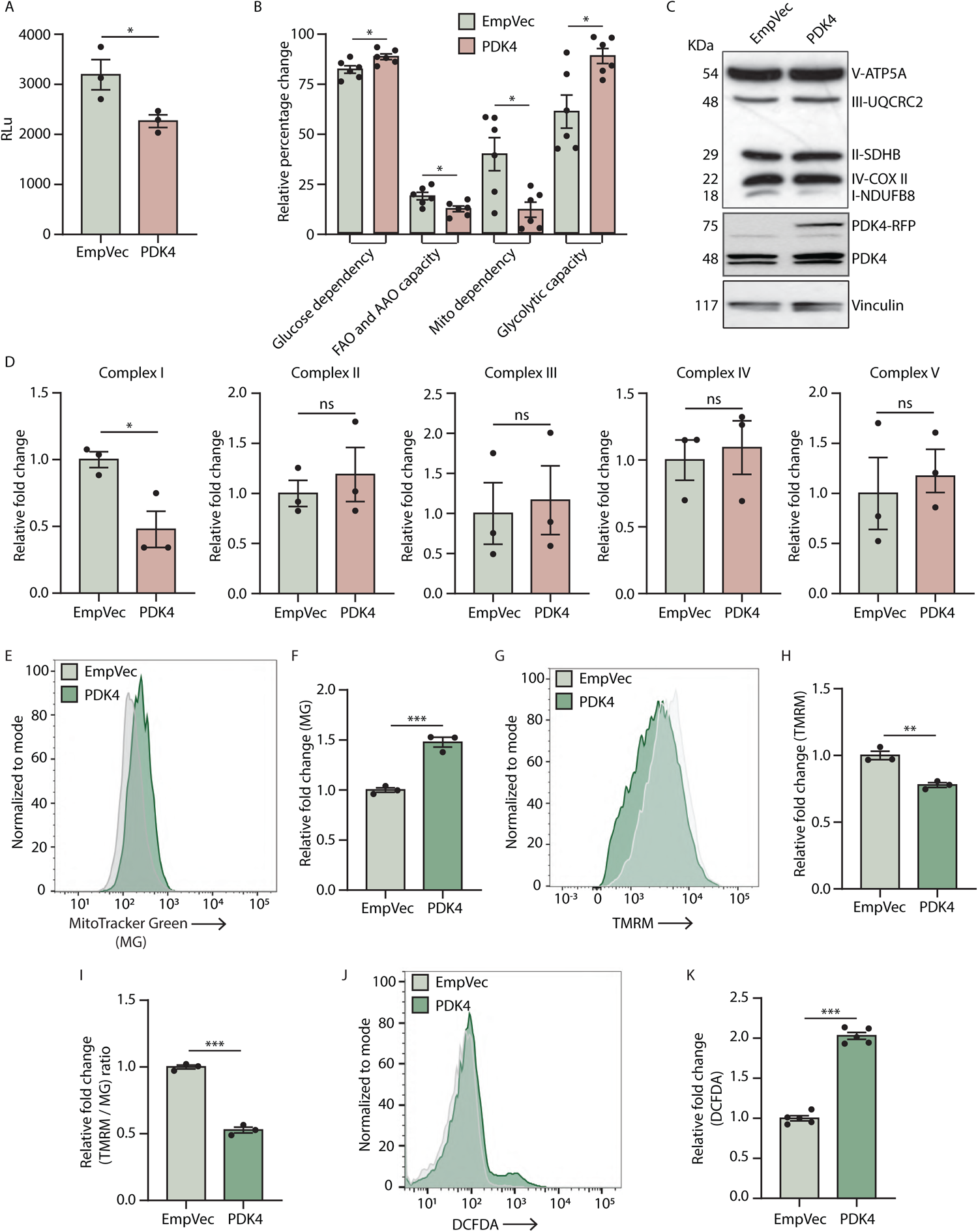
Increased PDK4 regulating mitochondrial functionality. (A) Graph represents total ATP production in SH-SY5Y cells transfected with empty vector (EmpVec) and PDK4, in RLu (relative luminescence) units. Data is shown as the mean ± SEM of 3 independent experiments. (B) Plot represents the percentage of ATP generated from glucose dependence, FAO (fatty acid oxidation) and AAO (amino acid oxidation) capacity, mitochondrial dependence and glycolytic capacity of cells transfected with indicated constructs. Data is shown as mean ± SEM of 3 independent experiments. (C) Cells transfected with indicated constructs were immunoblotted to detect expression of indicated proteins. (D) Graphs represent change in expression of different proteins, as shown in panel C. Data represents the mean ± SEM of 3 independent experiments. (E) Representative flow cytometry histogram plot shows the intensity of MitoTracker Green (250 nM) signals in empty vector (EmpVec) and PDK4 transfected cells. (F) Graph shows quantification of MitoTracker Green signals measured by FACS, as mentioned in E. Data is shown as mean ± SEM of 3 independent experiments. (G) Representative flow cytometry histogram plot shows the intensity of TMRM (250 nM) signals in empty vector (EmpVec) and PDK4 transfected cells. (H) Graph shows quantification of TMRM signals measured by FACS, as shown in panel G. Data is shown as mean ± SEM of 3 independent experiments. (I) Graph shows quantification of TMRM/MitoTracker Green signal ratios measured by FACS, as mentioned in panels E and G. Data is shown as mean ± SEM of 3 independent experiments. (J) Representative flow cytometry histogram plot shows the intensity of DCFDA (1 μM) signals in empty vector (EmpVec) and PDK4 transfected cells. (K) Graph shows quantification of DCFDA signals measured by FACS, as shown in panel J. Data is shown as mean ± SEM of 5 independent experiments. *\*p*≤0.05; *\*\*p*≤0.01; ns: not significant (estimated via unpaired two-tailed Student’s t-test).

### Excess PDK4 impairs autophagy

The stressed and dysfunctional mitochondria are known to be cleared by mitophagy ^50^. Moreover, ER-mito contact sites or MAM junctions can facilitate autophagosome formation ^19^. Further, compromised autophagy and mitophagy are known to be associated with multiple neurodegenerative diseases, including AD ^51^. LC3II/I ratio and P62 proteins were significantly higher in cells with exogenous expression of PDK4, suggesting impaired autophagic clearance (Figures 7A-7C). Live cell imaging using dual tagged mCherry-EGFP-LC3B showed significantly lower percentage of red puncta (marking acidic vesicles), compared to the controls (Figures 7D, 7E), supporting impaired autophagosomal-lysosomal clearance upon PDK4 over-expression. Next, we checked if compromised autophagy can affect the clearance of dysfunctional mitochondria. While, live cell imaging of cells co-transfected with control vector or PDK4 along with mito-RFP-GFP did not show any significant alteration across samples (Supplementary Figures 5A, 5B), treatment with 10μM CCCP for 1 hr elicited a drastic change in mitophagic clearance of dysfunctional mitochondria (Figures 7F, 7G). Significantly low numbers of red puncta marking mitochondria in lysosomes recapitulates our data suggesting blocked autophagosomal-lysosomal fusion ultimately culminating in compromised clearance of stressed mitochondria.

**Figure 7:**
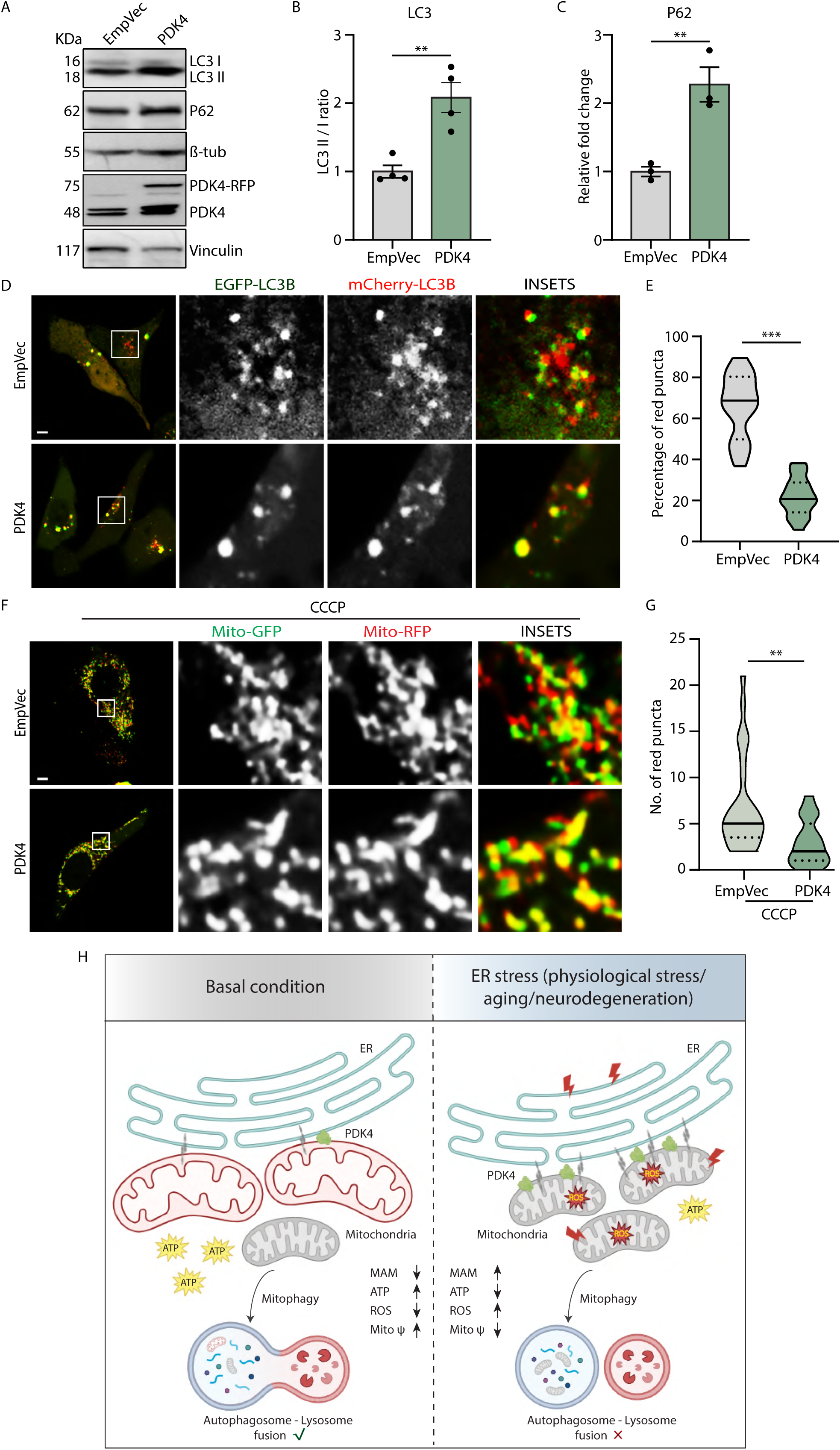
PDK4 over-expression regulates autophagic flux clearance. (A) SH-SY5Y cell lysates from empty vector (EmpVec) and PDK4 transfected samples were immunoblotted to detect expression of indicated proteins. (B) Graph represents change in ratio of LC3-II and LC3-I expression, as mentioned in A. Data represent the mean ± SEM of 4 independent experiments. (C) Graph represent change in expression of indicated protein, as mentioned in A. Data represents the mean ± SEM of 3 independent experiments. (D) Cells were transfected with mCherry-EGFP-LC3B, PDK4 and control (EmpVec), imaged under live-cell conditions. Enlarged views of the areas within the white boxes are shown (insets). Scale bar: 5 µm. (E) Violin plot shows the percentage of mCherry-LC3B (red) puncta (red puncta/total puncta*100), as described in D. Solid and dashed black lines mark median and quartiles respectively. Data shown are of ∼50 cells from 3 independent experiments. (F) Cells transfected with Mito-RFP-GFP, PDK4 and control (EmpVec), followed by treatment with CCCP (10 μM) imaged under live-cell conditions. Enlarged views of the areas within the white boxes are shown (insets). Scale bar: 5 µm. (G) Violin plot shows the Mito-RFP (red) puncta count, as described in F. Solid and dashed black lines mark median and quartiles respectively. Data shown are of ∼60 cells from 3 independent experiments. (H) A graphical representation demonstrating the impact of increased PDK4 in maintaining ER-mitochondrial communication and mitochondrial homeostasis. In basal condition, PDK4 level is relatively low, which corresponds to increased mitochondrial ATP production, membrane potential, lower ROS level and also lesser number of ER-Mito contact sites (MAM); Autophagosome-lysosome fusion remains functional. However, ER stress, aging and neurodegeneration-induced over-expression of PDK4 leads to altered mitochondrial homeostasis (more fragmented and depolarized mitochondria) and ER-mitochondrial communication (increased MAMs); reduced ATP production, increased ROS level, impaired autophagosome-lysosome fusion. Subsequently, exerting stress mediated disease progression. *\*p*≤0.05; *\*\*p*≤0.01; ns: not significant (estimated via unpaired two-tailed Student’s t-test).

## Discussion

In this study, we not only propose a non-canonical role of PDK4 in affecting mitochondrial morphology, function and clearance, we identify this regulator of oxidative phosphorylation as one of the connecting links between ER^UPR^ and mito^UPR^. Comprehensive analyses under various chemical as well as physiological ER stress conditions reveal PDK4 as a novel marker that gets deregulated at the mitochondria, and can hence act as a contributing factor to aging and neurodegeneration. We show that ER stress mediated elevated PDK4 levels affect ER-mitochondria contact sites, morphologically and functionally compromise mitochondria – factors essentially contributing to mito^UPR^ (Figure 7H).

There are distinct organellar mechanisms for coping with cellular stress at the ER and mitochondria, termed as ER^UPR^ and mito^UPR^, respectively. These primarily regulate accumulation of misfolded protein within the respective compartments. Build-up of unfolded, misfolded, or erroneously modified proteins in the ER lumen triggers a multilayered cellular response, activating one or more of the three canonical pathways of ER stress (IRE1α, PERK and ATF6). These pathways work in a well-coordinated manner to elicit increased chaperone expression to enhance protein folding, reducing protein load by translational attenuation and initiating ER-associated degradation (ERAD) process to get rid of the terminally misfolded proteins ^52^. Similarly, mito^UPR^ controls the response towards impaired mitochondrial protein import or accumulation of proteins in the matrix, leading to aberrant mitochondrial function. This stress response helps to restore mitochondrial homeostasis by activating chaperones like HSPD1 (Heat Shock Protein Family D (Hsp60) Member 1), HSPE1 (Heat Shock Protein Family E (Hsp10) Member 1) and proteases, including LONP1 (Lon protease 1); and can also induce mitochondrial fission by increasing DRP1 expression. Similar to ERAD, defective parts of mitochondria or the entire organelle may be eliminated by various degradation pathways, including, mitophagy, reticulo-mito-phagy, and mitochondria-associated degradation (MAD) ^20,50,53,54^. Our results posit PDK4 at a junction where it gets similarly deregulated by ER and mitochondrial stresses.

Many neurodegenerative diseases are characterized by the presence of misfolded protein aggregates, and upregulation of multiple ER stress markers ^55–58^. It is well established that ER is an extremely dynamic organelle, making contact crosstalk with various other cellular compartments at all times ^59–62^. Hence, it is logical to hypothesize that any cellular insult that elicits ER stress should also percolate to other organelles, simultaneously destabilizing them. While the complexity of organellar interconnectivity is progressively being unraveled, our knowledge of the molecular executors that sense and transmit ER stress to other organelles (probably as a defense mechanism to alleviate it) still remains sparse. Literature suggests increased mitochondrial fragmentation, abnormal functioning, cytosolic extrusion of mtDNA, impaired mitophagy – all features of deregulated mitochondria – are often associated with chronic neurodegeneration ^16, 63–65^. Further, ER-mito contact sites take part to regulate lipid, glucose, fatty acid-metabolism, calcium signaling, autophagosome formation, apoptosis; which are frequently found to be erroneous in neurodegeneration ^19, 66–70^. So, it is crucial to identify proteins from other organelles, especially mitochondria that would participate in organellar crosstalk during ER stress mediated neuronal death. Our results clearly suggest that PDK4 emerges as a potential mitochondrial sensor or effector of ER stress – future experiments, beyond the scope of this study would be required to fully establish this.

Studies have shown that Aβ peptide, a hallmark of AD may be generated at MAM junctions. Further, other essential molecular players, like amyloid-β protein precursor (AβPP), β-site APP cleaving enzyme 1 (BACE1), C-terminal fragment (CTF) of 99 amino acids (C99), the γ-secretase complex, and APP intracellular domain (AICD), closely associated with the pathogenesis of AD - are all detected at MAM junctions ^43, 71^. Aβ oligomeric aggregations are reported in the ER of hippocampal neurons, possibly contributing to their accelerated loss during neurodegeneration ^72^. Aβ plaque formation in mitochondria may lead to mitochondrial damage ^73^. In addition, it has been reported that MAM resident proteins like MFN2, sigma-1 receptor deficiency can lead to decreased ER-mito tethering, thereby reducing the deposition of toxic Aβ oligomers ^74, 75^. These evidences support the significance of MAMs in AD pathogenesis. Other neurodegenerative diseases like Parkinson’s disease (PD), Huntington’s disease (HD), Amyotrophic lateral sclerosis (ALS), Charcot-Marie-Tooth (CMT) disease, Wolfram syndrome (WS) are also associated with altered MAM formation or function ^76–80^. MAMs have been also suggested to be deregulated in various cancers ^81,82^, viral infection ^83^ and insulin resistance and diabetes ^84^. Interestingly, presence of PDK4 in MAM junctions is reported during aberrant insulin signaling ^29^. Together these support our findings that PDK4 gets recruited to ER-mito contacts during ER stress and its elevated level increase MAM formation and disturb mitochondrial homeostasis. Some features of neurodegeneration are also shared during the natural process of aging, one such being ER stress and consequential evocation of ER^UPR^ ^85^. Increased PDK4 protein levels in hippocampal region of aged rats as well as AD mice models clearly support this hypothesis. It is well documented that while UPR might have an initial neuroprotective role, sustained stress can initiate or potentiate the progress of neurodegeneration ^58^.

Altered mitochondrial dynamics is implicated in neurodegenerative diseases, including AD ^16, 86^. Imbalance in mitochondrial fission-fusion machinery can lead to abnormal neuronal function ^87^. Aβ driven post-translational modification of DRP1 (S-nitrosylation) is known to trigger mitochondrial fission, neuronal damage in AD ^14^. Here, we show PDK4 over-expression can lead to mitochondrial fragmentation; it is plausible to suggest that the PDK4-SEPT2-DRP1 axis could be utilized for this ^46^. However, detailed investigation of this is beyond the scope of the present study.

Brain is one of the highest energy-demanding organs; it is sensitive to even mild deficit in energy metabolism process. In AD, various players of the glycolytic pathway, TCA cycle, oxidative phosphorylation ^88^, specifically complex I of OXPHOS is significantly reduced ^89^. Our results show direct correlation between AD and higher levels of PDK4 in specific brain regions. Further, antagonistic effect of increased PDK4 on mitochondrial ATP production and functionality suggests that this could be one of the contributing factors in this neurodegenerative disease. We have also observed increased mitochondrial mass and ROS production in PDK4 over-expressed cells, which may generate an increased pool of damaged mitochondria (as modelled in ^90^). Increased mitochondrial mass is indicative of elevated oxidative stress, which is often reported in AD ^91, 92^. Neurodegenerative diseases are also often associated with deregulated autophagic clearance ^93^. Our results suggest attenuated fusion of autophagosomes with lysosomes upon exogenous expression of PDK4. More importantly, this drastically affects mitophagic clearance of dysfunctional mitochondria. There are evidences to suggest the involvement of PDK4 in regulation of autophagic flux under cellular stress; while it is reported that higher PDK4 levels can elevate autophagy ^94^, it is also shown to impair autophagy in VSMCs (Vascular smooth muscle cells) calcification ^95^. In our system, we find that while autophagy is induced upon PDK4 over-expression, autophagic degradation or the flux clearance is impaired. A mechanistic insight into this is being pursued as a separate study. PDK4 emerges as one of the potential mitochondrial targets that are altered during ER stress, a molecular phenomenon central to aging and neurodegeneration. Not only does ER stress affect PDK4 levels, this protein in turn can also positively regulate ER stress, while simultaneously deregulating mitochondrial morphology and function. This provides one of the direct evidences of ER^UPR^ regulating mito^UPR^.

## Material and Methods

### Constructs

Full-length PDK4 in pcDNA 4.1 and pDsRed1-N1 vector was generated. ER–GFP (GFP– KDEL) and CytERM–BFP were gifts from Erik L. Snapp (Janelia, USA). Wild type prion, prion mutant constructs (A117V, KHII) were gifts from Ramanujan S. Hegde (MRC-LMB, UK). mCherry-EGFP-LC3B was a gift from Terje Johansen (Arctic University, Norway). Mito-RFP-GFP was a gift from Benu Brata Das (IACS, India). Amyloid precursor protein intracellular domain (AICD-GFP) construct was a gift from Debashis Mukhopadhyay (SINP, India).

### Antibodies

Antibodies that were used are listed below:

| Antibody | Manufacturer (Cat no.) | Dilution |
| --- | --- | --- |
| *1.* PDK4 | Abcam, #110336 | 1:1000 |
| 2. BiP | Cell Signaling Technology (CST), #3177 | 1:1000 |
| 3. CHOP | Cell Signaling Technology (CST), #2895 | 1:1000 |
| 4. HERPUD1 | Invitrogen, #26301 | 1:1000 |
| 5. p-PDHA1(ser293) | Abcam, # 92696 | 1:2000 |
| 6. PDHA1 | Abcam, # 110334 | 1:2000 |
| 7. PDK1 | ABclonal, #A0834 | 1:1000 |
| 8. PDK2 | ABclonal, #A4737 | 1:1000 |
| 9. PDK3 | ABclonal, #A12480 | 1:500 |
| 10. MFN2 | Cell Signaling Technology (CST), #(D1E9),<br>11925 | 1:1000 |
| 11. VDAC1 | Abcam, #15895 | 1:1000 |
| 12. Calnexin | Cell Signaling Technology (CST),<br>#(C5C9),2679 | 1:1000 |
| 13. OXPHOS cocktail | Abcam, #110411 | 1:1000 |
| 14. LC3 | Novus Biologicals, #NB100-2220 | 1:1000 |
| 15. P62 | Novus Biologicals, #NBP1-48320 | 1:2000 |
| 16. ATF5 | Abcam, #184923 | 1:1000 |
| 17. Vinculin | Abcam, #129002 | 1:5000 |
| 18. β-actin | Abcam, # 8226 | 1:5000 |
| 19. GAPDH | Santa Cruz Biotechnology (SCBT), #SC365062 | 1:5000 |
| 20. Anti-rabbit IgG, HRP-linked Antibody | Cell Signaling Technology (CST), #7074 | 1:3000 |
| 21. Anti-mouse IgG, HRP-linked Antibody | Cell Signaling Technology (CST), #7076 | 1:3000 |
| 22. RFP | Ramanujan S. Hegde (MRC-LMB, UK) | 1:5000 |

### Reagents

Thapsigargin (T9033), Tunicamycin (T776), Lipofectamine 2000 (Thermo Fisher Scientific, #11668019), MitoTracker Red FM (Thermo Fisher Scientific, #22425), MitoTracker Deep Red FM (Thermo Fisher Scientific, #22426), MitoTracker Green FM (Thermo Fisher Scientific, #M7514), CM-H2DCFDA (Thermo Fisher Scientific, #C6827), TMRM (Thermo Fisher Scientific, #T668), CCCP (Sigma-Aldrich, #C2759), percoll (Sigma-Aldrich, #P1644), Amyloid β Protein Fragment 1-42 (Aβ42) (Sigma-Aldrich, #A9810), Luminescent ATP detection assay kit (Abcam, #113849), 2-Deoxy-D-glucose (2-DG) (TCI, #154-17-6), Oligomycin A (Sigma-Aldrich, #75351).

### Animal experiment

Age matched *Apoe* knock-out C57BL/6J mice and young and old Sprague-Dawley male rats were bred and housed under SPF conditions in the Institutional Animal Facility of CSIR-IICB and different brain region specific lysates were provided by Arun Bandyopadhyay and Joy Chakraborty (CSIR-IICB).

### Human tissue sample

All human brain lysates (control, AD patients) were obtained from Novus Biologicals, LLC (NB820-59177, NB820-59363).

### Cell culture

SH-SY5Y Human neuroblastoma cells were a gift from Debashis Mukhopadhyay (SINP, Kolkata, India). Cells were grown in 10% fetal bovine serum (FBS - HI; Gibco) in Dulbecco’s modified Eagle’s medium (DMEM), (Gibco, #12100046) at 37°C and 5% CO2. For transfections of cells, Lipofectamine 2000 was used (Invitrogen, Carlsbad, CA, USA) as per the manufacturer’s instructions. 24 h after transfection, cells were lysed using suitable lysis buffer.

### Immunocytochemistry

For immunocytochemistry, cells were fixed with either 4% formaldehyde or methanol as per the requirement of the antibody, as described previously (Chakrabarti and Hegde, 2009; Srivastava and Chakrabarti, 2014). Cells were permeabilized using phosphate-buffered saline (PBS) containing 10% FBS and 0.1% saponin (Sigma-Aldrich) for 60 min, followed by overnight staining in primary antibody at 4°C and 60 min incubation in secondary antibody at room temperature. The samples were then imaged using confocal microscope.

### Confocal imaging and image analyses

Confocal imaging was performed using LSM 980, LSM710 ConfoCor 3 and Nikon A1R+Ti-E with N-SIM and FCS microscope systems. A diode laser (BFP excitation with 405nm line), an Ar-ion laser (GFP, Alexa Fluor 488 excitation with the 488 nm line) and He-Ne lasers (RFP, Alexa Fluor 546 excitation with the 543 nm line; MitoTracker Red excitation with the 561 nm line; Alexa Fluor 633 and MitoTracker Deep Red excitation with 640 nm line) were used with 63×1.4 NA oil immersion objectives. Transfected cells were imaged in CO_2_-independent medium (Thermo Fisher Scientific), maintaining conditions of live-cell imaging as described previously (Mitra and Lippincott-Schwartz, 2010).

Mitochondrial length measurement was done manually by drawing lines along the entire length of each mitochondrion from 2D confocal micrographs of cells of resolution 1024×1024 at 300 dpi; where mitochondria were stained with MitoTracker Red. mCherry-LC3B and Mito-RFP puncta were counted from 2D confocal micrographs of cells of resolution 1024×1024 at 300 dpi. Measurements were done using Fiji. Co-localization analysis of ER-mitochondria and ER-mitochondria-PDK4 signals were carried out using MATLAB image processing tool and Fiji.

### Western blotting

The protocol for western blotting was as described previously (Kaul and Chakrabarti, 2017). 10% Tris-tricine gels or 7.5% Tris-glycine gels were used for SDS–PAGE followed by western blotting. Quantification of western blots was performed using GelQuant (Thermo Fisher Scientific) software. At least three independent experiments were performed, and band intensities were normalized to loading controls. p-values were determined using Student’s t-test.

### Isolation of MAM, Mitochondria, ER and Cytosolic fractions

The MAM fraction was isolated following an established protocol ^96^. Briefly, cells were cultured in 100mm plates, followed by either TG or vehicle treatment, they were washed with ice-cold PBS, collected with cell scraper and kept on ice. Cells were centrifuged and the pellet was re-suspended in homogenization buffer (30mM Tris-HCl (Ph 7.4) buffer containing 225mM mannitol, 75mM sucrose and protease inhibitors). Homogenization procedure was performed manually with 20 strokes of a 27-gauge needle, on ice. At first, nuclei and unlysed cells were pelleted by centrifugation at 600g for 5min. This was repeated and after the second centrifugation the supernatant was collected. This was followed by a spin at 7000g for 10min. The pellet was further processed for obtaining pure mitochondria and MAM fraction. The supernatant was further centrifuged at 20,000g for 30min. Enriched ER and cytosolic fractions were separated in the pellet and supernatant respectively, when the collected supernatant was subjected to ultra-centrifugation at 1,00,000g for 1h. The pellet containing mitochondrial fraction was re-suspended in Homogenization buffer, this was centrifuged then at 7000g for 10min and again after re-suspension of the pellet in the same buffer, centrifugation of 10,000g for 10min was done. The collected pellet was re-suspended in MRB buffer (250mM mannitol, 25mM HEPES (pH 7.4) and 1mM EGTA) and layered on top of 30% percoll medium (225mM mannitol, 5mM HEPES (pH 7.4) and 0.5mM EGTA) and centrifuged at 95,000× *g* for 30 min. The MAM fraction was extracted from percoll gradient by centrifugation at 6300g at 10min and further purified by ultra-centrifugation at 1,00,000g for 1h to remove remaining mitochondrial fraction. Similarly, the pure mitochondria fraction was collected from percoll gradient and centrifuge at 6300g for 10min to obtain mitochondrial pellet. The entire fractionation procedure was performed throughout at 4°C. Fractions were analyzed by Western blotting with suitable antibodies for different fractions.

### Quantitative real-time PCR (qRT-PCR)

RNA extraction from SH-SY5Y cells is carried out using TRIzol reagent (Thermo Fisher Scientific) for control and drug treated or transfected samples. cDNA synthesis from total RNA is performed using High-Capacity cDNA Reverse Transcriptase Kit (Applied Biosystems) following the manufacturer’s protocol.

Quantitative real-time experiments were performed using PowerUP SYBR Green Master Mix (Applied Biosystems, #A25742) with gene specific primers, according to the manufacturer’s instructions. The primer sequences of the related genes are – PDK4 (forward primer, 5′-AGATACACTCATCAAAGTTCGAA-3′; reverse primer, 5′-TCATCAGCATCCGAGTAGA-3′), BiP (forward primer, 5′-GAAAGAAGGTTACCCATGCAGT-3′; reverse primer, 5′-CAGGCCATAAGCAATAGCAGC-3′), CHOP (forward primer, 5′-CCAGCAGAGGTCACAAGCAC-3′; reverse primer, 5′-TTCTCCTTCATGCGCTGC-3′), XBP1-s (forward primer, 5′-GAGTCCGCAGCAGGTGC-3′; reverse primer, 5′-TGATGACGTCCCCACTGAC-3′), β-Actin (forward primer, 5′-CACTGGCATCGTGATGGA-3′; reverse primer, 5′-CCGTGGCCATCTCTTGCT-3′). ND2, SDHA, CYTB, COX-II and ATP8 primers were gift from Soumen Kanti Manna (SINP, Kolkata, India). β-Actin was used as the reference gene. Relative expression of each gene was measured using the gene’s 2^-ΔΔCt^.

### mRNA Transcriptomics

RNA was isolated as described above. The quantity and quality of the isolated RNA were evaluated using Qubit 4.0 (Thermo Fisher Scientific). Transcriptome sequencing was performed using Illumina Novaseq 6000 and has been deposited in NCBI’s Gene Expression Omnibus ^97^ and are accessible through GEO Series accession number GSE266545 (https://www.ncbi.nlm.nih.gov/geo/query/acc.cgi?acc=GSE266545). A fold change of ±2 in the log2 scale and p-value of 0.01 were set as a cutoff for the categorization of Differentially Expressed Genes (DEGs). MitoCarta database (v2.0) was used to identify mitochondrial genes which were among the DEGs.

### Multiple sequence alignment (MSA)

Full length FASTA sequences of proteins were retrieved from NCBI protein (https://www.ncbi.nlm.nih.gov/protein) database and provided as input in Clustal Omega (https://www.ebi.ac.uk/jdispatcher/msa/clustalo). Sequence identity matrix was obtained from the webserver.

### ATP measurement

The method for ATP measurement was carried out as described previously ^98^. Cells were counted and 1×10^5^ cells/ml were seeded for each sample. Transfection was performed next followed by treatment with DMSO, 2-Deoxy-D-Glucose (DG, final concentration 100mM), Oligomycin A (Oligo, final concentration 1μM), or combined DG and Oligomycin A for 60min. ATP production was measured with Luminescent ATP detection assay kit (Abcam, #113849), following manufacturer’s protocol. After cell lysis and substrate addition, luminescence was detected using BioTek SYNERGY HTX Reader at 540 nm wavelength.

### ROS production analysis

ROS generation was measured using DCFDA (final conc.1μM). In brief, cells were washed with PBS, stained, incubated at 37 °C for 30 min, washed with PBS and measured using a flow cytometer (BD FACSAria II) and analysed using FlowJo Software (v10.9.0).

### Mitochondrial potential and mass measurement

Mitochondrial membrane potential and mass were measured using TMRM and MitoTracker Green co-stained together (final conc. 250nM) at 37 °C for 30 min, then washed with PBS and measured using a flow cytometer (BD LSRFortessa) and analysed using FlowJo Software (v10.9.0).

### Statistical analysis

Data are expressed as mean ± SEM. Statistical comparisons were performed using a paired Student’s t test and one way ANOVA whichever is applicable. Differences were considered statistically significant when the p-value was <0.05.

## Supporting information

All Supplementary Figures merged

## Acknowledgments

We thank Erik L. Snapp (Janelia, USA), Ramanujan S. Hegde (MRC-LMB, UK), Terje Johansen (Arctic University, Norway), Benu Brata Das (IACS, India) for constructs; Debashis Mukhopadhyay (SINP, India) for cells and construct. We thank Shilpak Chatterjee (IICB, India) and Anwesha Kar for helping with flowcytometric data analysis; Subhrangshu Das for image analysis; Soumen Kanti Manna for primers; Sk. Ramiz Islam for helping with ATP luminescence study and Palamou Das for providing AICD transfected (Aβ42 treated) samples. We acknowledge all past and present members of S.C. and O.C. laboratories, for their support throughout the study, especially Sarpita Bose.

PM and SC are supported by the intramural funding of CSIR - Indian Institute of Chemical Biology. SM and OC are supported by the intramural funding of the Department of Atomic Energy (DAE), Government of India. OC is partially funded by SERB, Department of Science & Technology (CRG/2021/000638), Government of India and National Women Bioscientist Award grant, Department of Biotechnology, Government of India. PM acknowledges UGC (20/12/2015(ii)EU-V, #2121530853) for PhD fellowship. SM acknowledges CSIR (09/489(0109)/2018-EMR-I) for PhD fellowship. RM and TR acknowledge the Council of Scientific and Industrial Research (CSIR), India for PhD fellowship.

## Author contributions

PM, SM, OC and SC conceived the project and designed the experiments. PM and SM performed the experiments and analyzed the data. RM and TR helped with the animal studies. PM, OC and SC interpreted the results. PM, OC and SC wrote the manuscript and coordinated the project. All authors contributed to the article, approved the submitted version.

