## Supplementary material for "The impact of ER^UPR^ on mitochondrial integrity mediated by PDK4": All Supplementary Figures merged

### Supplemental information titles and legends

**Supplementary Figure 1: Effect of ER stress inducing drugs on cellular homeostasis**

(A) Immunoblotting of SH-SY5Y cell lysates was carried out to check indicated protein levels in TG (0.5µM for 3 hrs) induced ER stress, compared to control (DMSO). (B) Graphs represent change in expression of different proteins, as mentioned in A. Data represent the mean ± SEM of 3-4 independent experiments. (C) Immunoblotting of cell lysates was done to detect indicated protein level in Tun (Tunicamycin, 5µg/ml for 3 hrs) induced ER stress, compared to control (DMSO). (D) Graph shows change in expression of indicated protein level, as mentioned in C. Data represent the mean ± SEM of 3 independent experiments. (E) Table shows sequence identity matrix (calculated in percentage) of indicated proteins, generated using the Clustal Omega software. *$p \leq 0.05$; **$p \leq 0.01$ (unpaired two-tailed Student's t-test).

**Supplementary Figure 2: Prion disease mutant alters PDK4 level**

(A) Immunoblotting of SH-SY5Y cell lysates, transfected with indicated constructs were subjected to immunoblotting for detecting protein expression of different proteins. (B) Graphs represent change in expression of different proteins, as mentioned in A. Data represent the mean ± SEM of 3 independent experiments. *$p \leq 0.05$; ns: not significant (one-way ANOVA with Bonferroni corrections).

**Supplementary Figure 3: Thapsigargin mediated alteration in mitochondrial morphology**

(A) Flowchart shows the sequential steps of isolating different cellular fractions from the whole cell lysate, i.e. nuclei, cytosol (cyto), mitochondria (mito), MAM and ER. SH-SY5Y cells were subjected to treatment with either DMSO or TG (0.5µM for 3 hrs) prior to the fractionation process. (B) Cells treated with TG or DMSO, imaged under live-cell conditions usingMitoTracker Red FM. Enlarged views of the areas within the white boxes are shown (insets). Scale bar: 5 µm. (C) Violin plot of the mitochondrial length (µm) in images, as described in B. Solid and dashed black lines mark median and quartiles respectively. ***$p \leq 0.001$ (unpaired two-tailed Student's t-test).

**Supplementary Figure 4: Effect of PDK4 over-expression on OXPHOS RNA levels**

qRT-PCR analyses show non-significant changes of indicated OXPHOS mRNA levels, in control (EmpVec) and PDK4 transfected SH-SY5Y cells. Data represent the mean ± SEM of 3 independent experiments. Ns: not significant (unpaired two-tailed Student's t-test).

**Supplementary Figure 5: Effect of PDK4 over-expression on mitophagy**

(A) Cells transfected with Mito-RFP-GFP, PDK4 and control (EmpVec), imaged under live-cell conditions. Enlarged views of the areas within the white boxes are shown (insets). Scale bar: 5 µm. (B) Violin plot shows the Mito-RFP (red) puncta count, as described in A. Solid and dashed black lines mark median and quartiles respectively. Data shown are of ~60 cells from 3 independent experiments. Ns: not significant (unpaired two-tailed Student's t-test).

Supplementary Figure 1

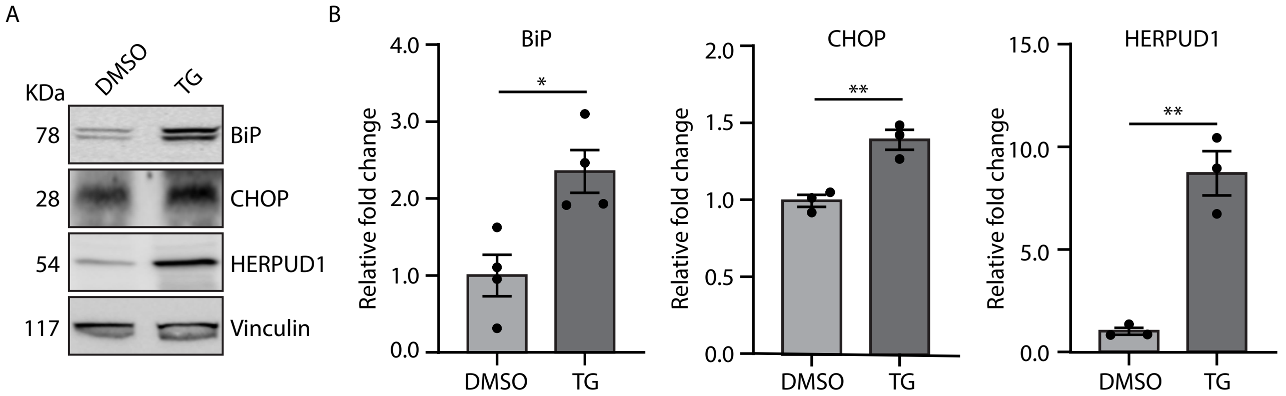


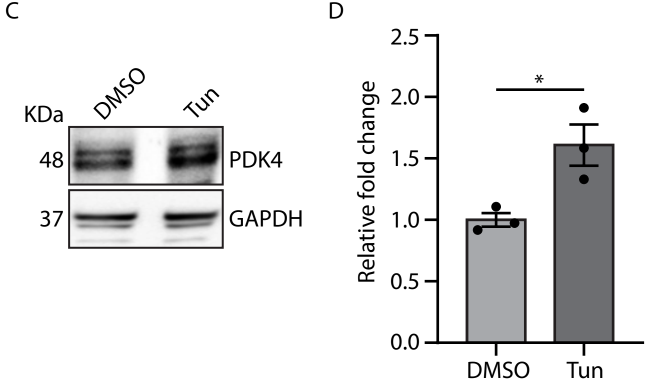


E

Sequence identity matrix (percentage)

| | PDK4 | PDK3 | PDK1 | PDK2 |
|---|---|---|---|---|
| PDK4 | 100.00 | 62.59 | 64.72 | 64.52 |
| PDK3 | 62.59 | 100.00 | 67.33 | 67.00 |
| PDK1 | 64.72 | 67.33 | 100.00 | 68.06 |
| PDK2 | 64.52 | 67.00 | 68.06 | 100.00 |

Supplementary Figure 2

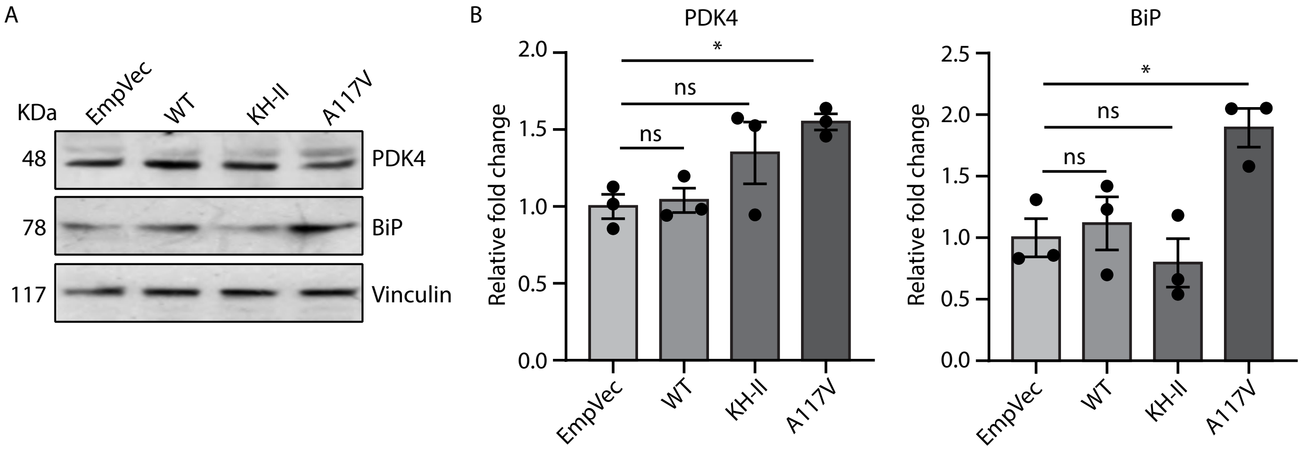

Supplementary Figure 3

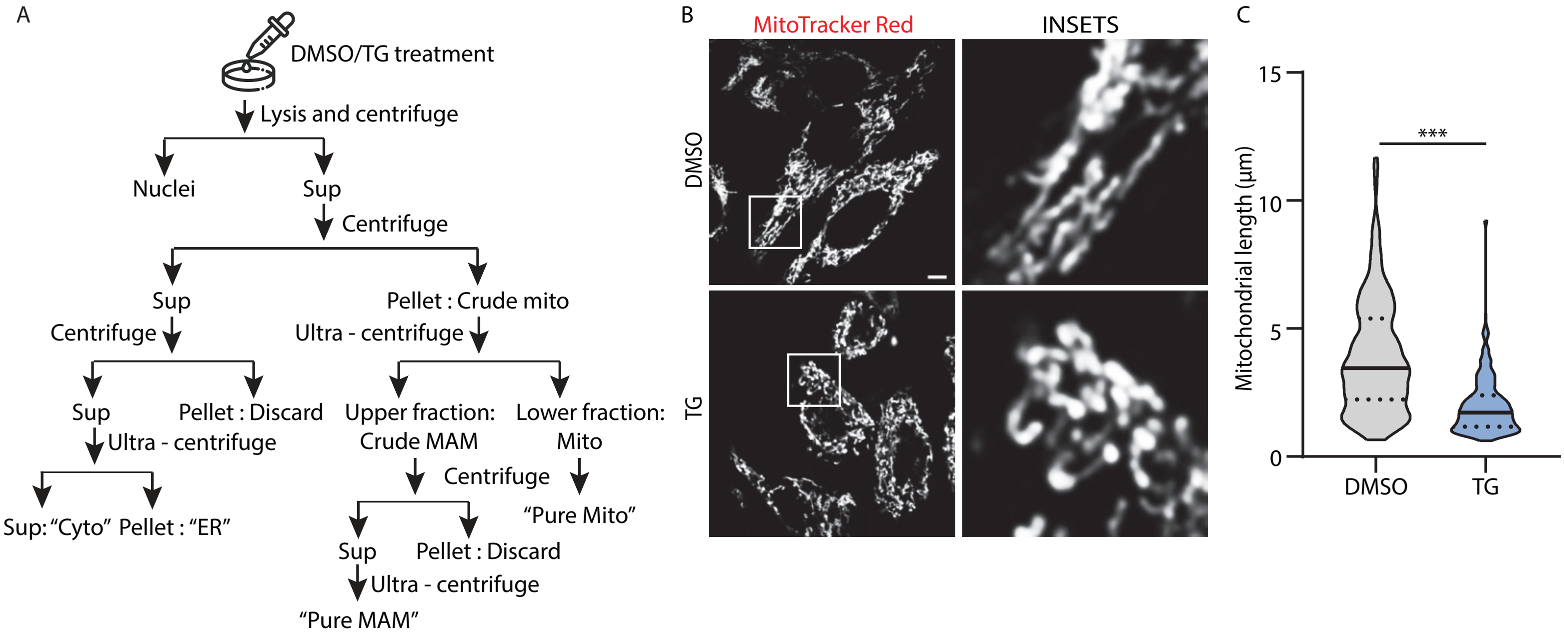

Supplementary Figure 4

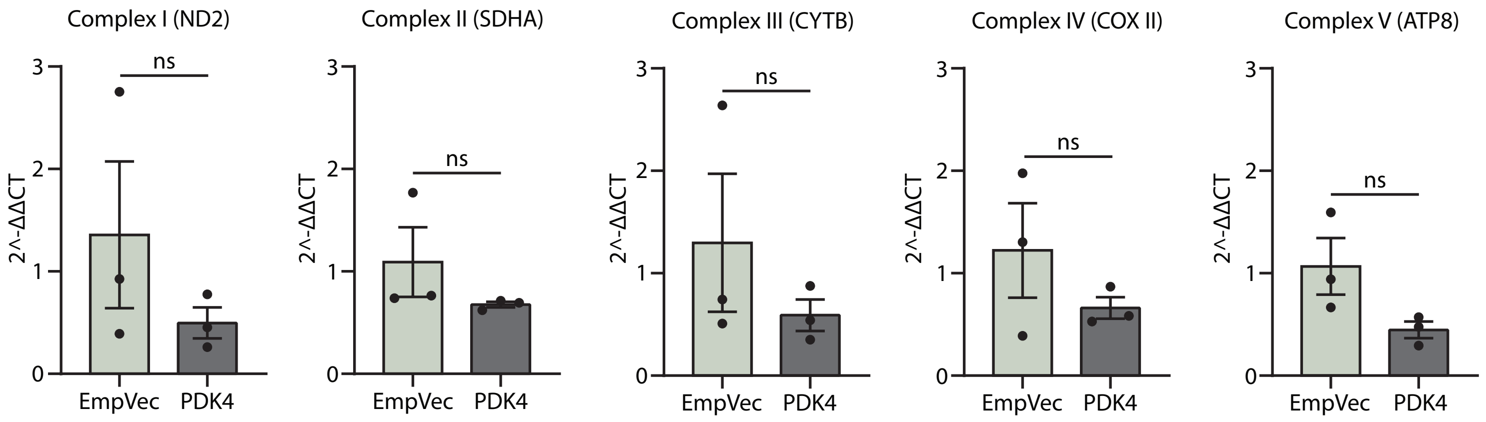

Supplementary Figure 5

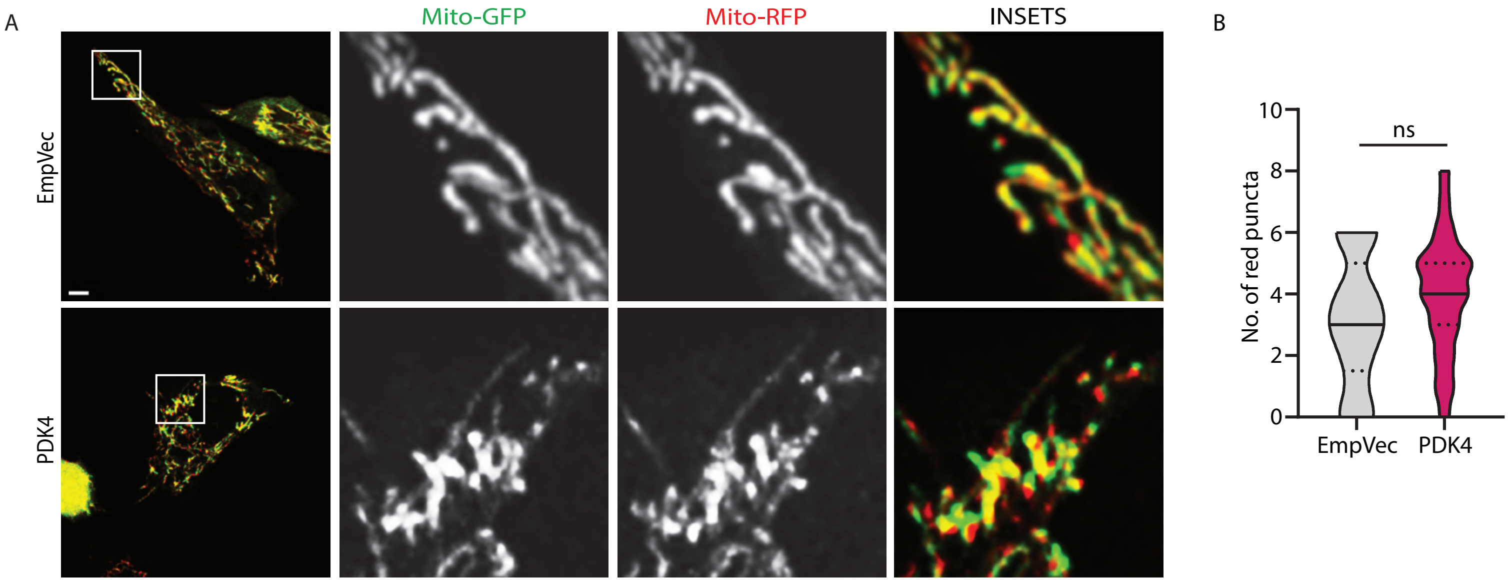
